# Multi-modal MRI characterisation of vascular remodelling, muscle fibre integrity, and inflammatory recovery in a hindlimb ischaemia mouse model

**DOI:** 10.64898/2026.07.17.739107

**Authors:** C.J. Lyons, M.N. Doulgkeroglou, C. Sanz-Nogués, XZ. Chen, C.A. Lagonda, T. O’Brien, N. Colgan

**Author notes:** Equal contribution (T. O’Brien, N. Colgan).

## Abstract

The hindlimb ischaemia (HLI) mouse model is a widely used preclinical model of chronic limb-threatening ischaemia (CLTI). While CLTI involves complex interactions between impaired perfusion, inflammation and muscle wasting, the standard imaging approach, laser Doppler imaging (LDI), only assesses perfusion. MRI is used clinically to assess neural tracts, inflammation, and perfusion in the brain. We therefore evaluated whether a multimodal MRI approach could longitudinally monitor recovery in the HLI mouse model. Mice underwent MRI three days pre-HLI surgery, and on Days +3 and +7 post-surgery, with histology on Day +7. The MRI detected significant increases in muscle volume and inflammation after HLI surgery, with significant decreases in perfusion, vascular length, and muscle fibre integrity. Overall, MRI can monitor inflammation, muscle fibre integrity, and vascular recovery post-HLI and should be applied in future studies to identify mechanisms of therapeutic recovery in a sequential in vivo analysis without requiring animal sacrifice.

Graphical Abstract

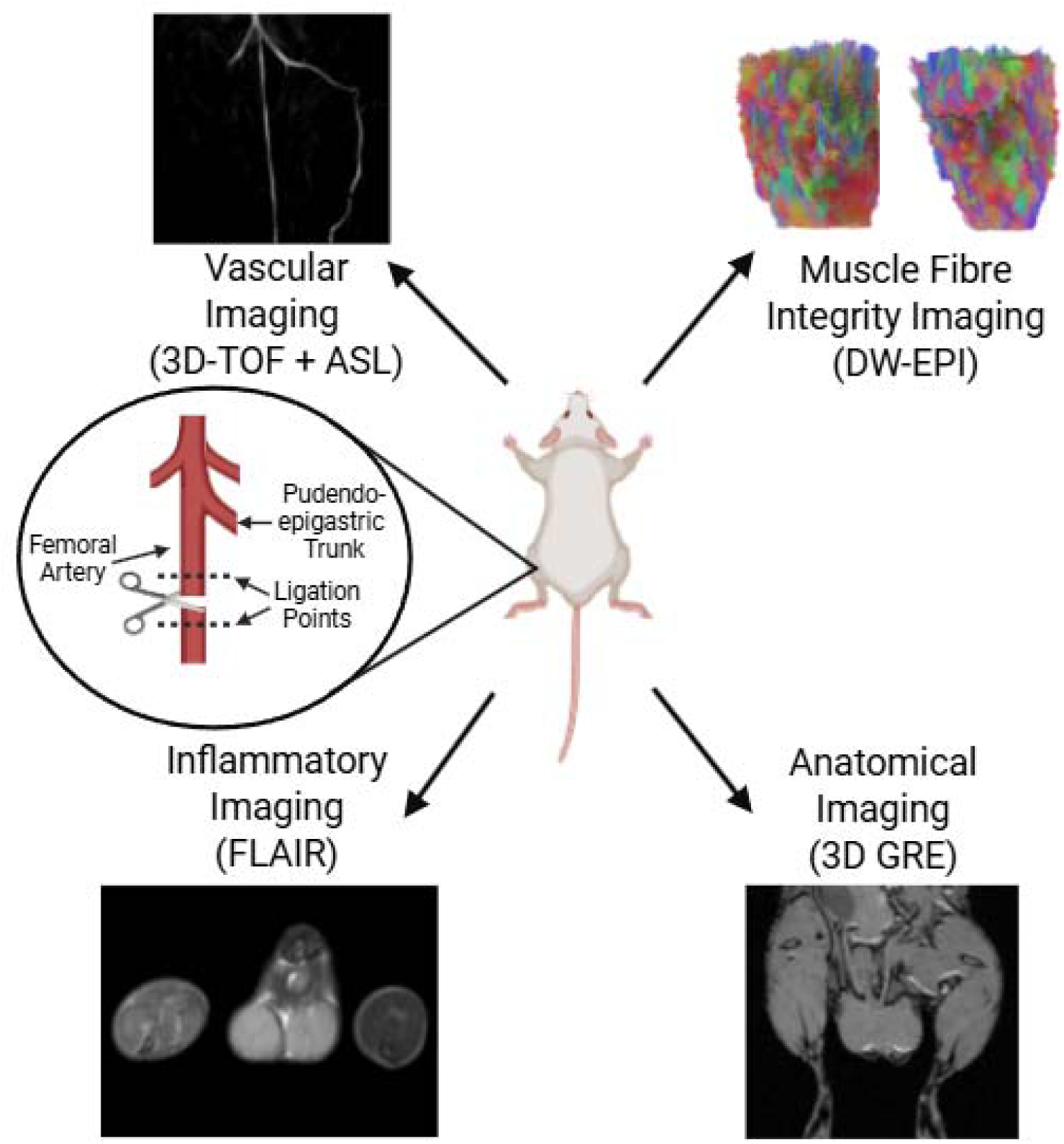

---

Chronic limb-threatening ischaemia (CLTI) is a multifaceted condition resulting from progressive atherosclerotic narrowing and occlusion of the lower limb arteries. The resulting reduction in blood flow creates a severely hypoxic and inflammatory tissue environment, leading to progressive tissue damage and loss of limb function [1]. CLTI presents as ischaemic rest pain, non-healing ulcers and/or gangrene in the affected lower limb. An estimated 10-25% of CLTI patients do not respond to standard revascularisation therapies and continue to progress to advanced disease, placing them at risk of major limb amputation [2,3]. Overall, CLTI carries a poor prognosis, with a 5-year mortality rate typically exceeding 50% [4,5]. Consequently, there is an urgent need to develop novel therapeutic strategies for this patient population.

The hindlimb ischaemia (HLI) mouse model is widely used as a preclinical model for evaluating potential therapeutics for CLTI, with laser Doppler imaging (LDI) being the most common imaging modality to longitudinally monitor blood flow recovery following HLI induction [6–9]. LDI provides high-resolution (0.1 mm) assessment of limb perfusion recovery but does not quantify the inflammatory response, muscle volumetric changes, or alterations in muscle architecture, all of which are substantially affected in CLTI and HLI. This is particularly relevant as many therapeutics being investigated for CLTI, including mesenchymal stromal cells, exert pleiotropic effects that extend beyond revascularisation, including promotion of muscle regeneration and the reduction of inflammation and fibrosis [10–12]. In most cases, endpoint histology is required to quantify these effects. Longitudinal assessment of these parameters in vivo would therefore require euthanasia and tissue collection at each timepoint, preventing serial evaluation within the same animal.

Magnetic resonance imaging (MRI) is routinely used clinically for the non-invasive monitoring of pathophysiology in soft tissues. Advances in MRI technology have enabled the assessment of pathophysiological changes in the HLI pre-clinical model; however, several groups have primarily focused on the use of MRI to monitor vascular recovery over time [13–17]. This study aims to integrate complementary MRI sequences and analytical techniques in a longitudinal multimodal imaging framework to investigate the HLI mouse model, enabling the simultaneous assessment of vascular remodelling, muscle integrity, and inflammatory recovery over time [13,18]. By combining these complementary analyses, MRI can characterise the temporal pathological cascade underlying the functional recovery and provide a comprehensive understanding of the trajectory of limb recovery after an ischaemic injury.

## Methodology

### Animals

Six male 7-8 week old BALB/cAnN-Foxn1 nu/Rj mice were sourced from Janvier Labs and were housed in a licensed preclinical facility at the University of Galway, with monitoring and support from qualified animal technicians and a veterinary surgeon. Ethical approval was granted by the Institutional Animal Care Research Ethics Committee. Project authorisation was granted by the Health Product Regulatory Authority (HPRA) in Ireland. Animals were then acclimatised for seven days prior to any procedure (Figure 1).

**Figure 1.**
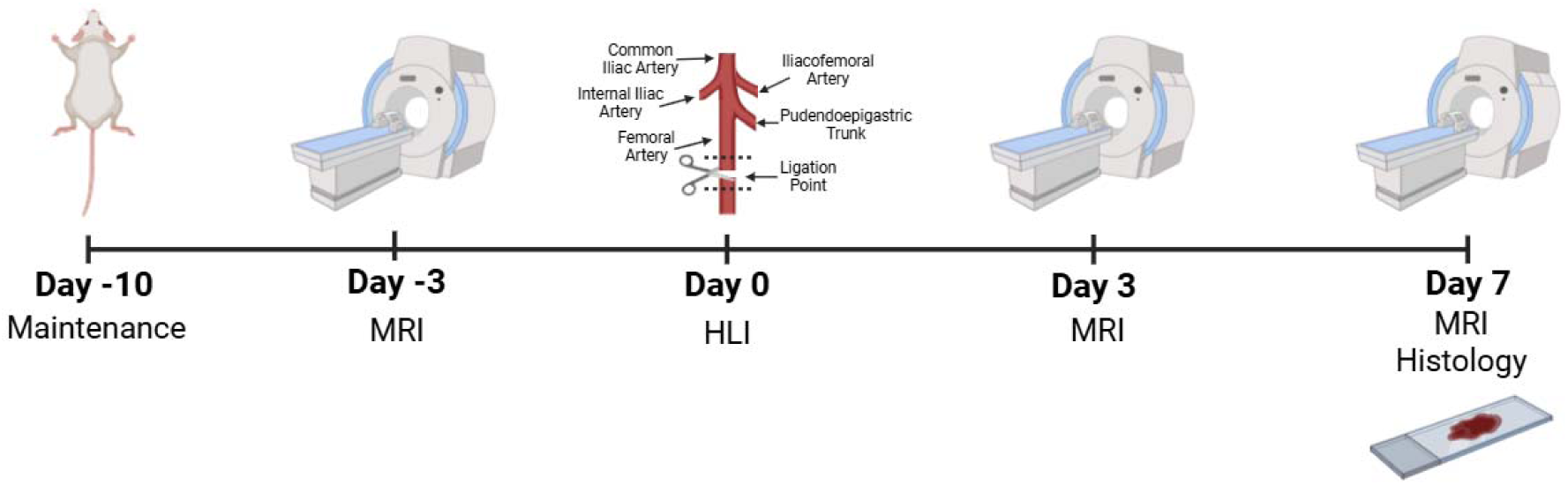
Study timeline. Scans were carried out on a 3T MRI system. MRI Scans include: DW-EPI (muscle fibre integrity), 3D GRE (muscle volume), FLAIR (inflammation), 3D-TOF (angiogenesis), and ASL (perfusion).

### Induction of HLI

On Day 0, animals were anaesthetised with 100 mg/kg of Ketamine and 0.5 mg/kg of Medetomidine injected subcutaneously prior to surgery. The HLI surgery consisted of unilateral ligation of the left femoral artery distal to the pudendoepigastric branch and then transecting the artery (Figure 1). Animals were administered with 0.1 mg/kg of buprenorphine thrice daily for pain management. Animals were then euthanised on Day +7 after surgery using CO_2_.

### MRI

Animals were scanned with a nanoScan 3T (Mediso Ltd) with a 42 mm Mouse WB RF coil and Nucline software (v. 3.05.12) on Day −3 before HLI surgery and then on Days +3 and +7 after surgery. Animals were anesthetised using 5% isoflurane and then maintained with 1.5-2% isoflurane in a temperature-controlled environment (37°C). Mice were placed in a prone position with their legs raised to facilitate the hindlimbs resting on the horizontal plane as noted in a paper by Bryant [18].

After performing a scout on each animal, a 3D Shim was carried out, focusing on the hindlimbs. Scans were then performed in the following order: 3D Multi-TOF, ASL, FLAIR, DW-EPI, and 3D-GRE, with a total scan time of ∼2 hrs. The settings for each scan are outlined below.

3D Multi-TOF: scan length = 7.2 mm, slice number = 36, resolution = 0.21 x 0.21 x 0.2 mm, number of fields of view = 5, overlap slices = 8, number of excitations = 2, TR = 25 ms, TE = 4.4 ms, flip angle = 30°.

FLAIR imaging: slices number = 30, slice gap = 0.1 mm, resolution 0.238 x 0.238 x 1.1 mm, excitation number = 4, TR = 8379 ms, TI = 4179 ms, TE = 33.1 ms, and an echo trail length = 8.

A flow-sensitive alternating inversion recovery sequence [19] echo planar imaging readout, ASL, was implemented with the following parameters: slice number = 12, resolution = 0.781 x 1.042 x 1 mm, excitation number = 10, TR = 3.6 ms, TE = 18 ms, flip angle = 30°, post labelling delay = 3204 ms, and an arterial spin labelling repetition time = 4909 ms.

DW-EPI: slice number = 15, resolution = 0.4 x 0.4 x 0.4 mm, slice gap = 0.4 mm, excitation number = 5, TR shot number = 1, TR = 3000 ms, TE = 26 ms, B value = 500 s/mm^2^ with three B = 0 s/mm^2^ images.

3D-GRE: resolution = 0.25 x 0.25 x 0.22 mm, slice number = 108, number of excitations = 4, TE = 6.3 ms, TR = 20 ms, flip angle = 10°.

### MRI Image Analysis

MRI muscle volumetric and FLAIR assessment was measured using 3D Slicer (v. 5.8.0) by creating volumes of interest around the calf muscles and calculating the enclosed volume and average pixel intensity, respectively. Representative images of each scan type were taken using InterView Fusion (v 3.11.010.000).

To calculate the water diffusion parameters MRIcroGL (v 1.2.20220720) was used to convert the DICOM file into a NIfTI format. This was then converted into .sz file format using DSI Studio (v. Hou “侯”). FSL Topup and FSL eddy correction were carried out using the NIfTI file with subsequent motion, orientation correction, and then b-table rotation. The restricted diffusion was quantified using restricted diffusion imaging, as per Yeh et al. [20]. The diffusion data were then reconstructed using generalized q-sampling imaging [21] and a diffusion sampling length ratio of 0.6.

To analyse the surface profile of the vasculature the 3D-TOF data were analysed using 3D Slicer coupled with the Vascular Modelling Toolkit extension [22]. 3D-TOF images were thresholded and then the Vascular Modelling Toolkit extension was used to identify the endpoints, and centreline, after which a centreline curve was generated. Afterwards, the longest combined length of artery was quantified for both left and right limbs. The longest combined length of artery was calculated by beginning at the descending aorta-common iliac branch point until the furthest arterial branch in each limb extremity as the endpoint.

Pulsed ASL was performed by using the proximal inversion with control for off-resonance effects technique (PICORE) as described by Buxton’s et al. [23]. The process is defined by three functions: (a) the delivery function c(t) is the normalized arterial concentration of magnetisation arriving at the voxel at time t; (b) the residue function r(t,t’) is the fraction of tagged water molecules that arrived at time t’ and are still in the voxel at time t; and (c) the magnetisation relaxation function m(t,t’) is the fraction of the original longitudinal magnetisation tag carried by the water molecules that arrived at time t’ that remains at time t. For the perfusion maps, it is assumed that the off-resonance and magnetisation transfer effects of the tag and control pulses are identical. In-house software implemented in MATLAB was used to process the images and generate peak perfusion maps. An ROI was placed on both the right and left limbs to measure the perfusion (ml/min/100g) in the hind limb.

### Histology

The calf muscles were carefully dissected from ischaemic and non-ischaemic limbs. The soleus was then further separated from the gastrocnemius and plantaris, and both set of muscles were imaged, and their wet weight recorded. Tissue samples were fixed with 10% buffered formalin for 48 hrs at room temperature and processed for histological analysis using the Epredia™ Excelsior™ AS Tissue Processor. Samples were then embedded in wax transversally, and 5 μm sections were taken. Sections were taken every 30 μm through the muscles with 5-6 sections per muscle analysed.

Sections were stained with Mayers Haematoxylin (Sigma, MHS32) and Eosin Y (Sigma, HT110232) (H&E) as follows. Slides were rehydrated with xylene (2 x 10 mins) then rehydrated via an ethanol series of 100%, 100%, 95%, 70%, and 50% and deionised water for 2 mins each. Samples were then stained with Mayers Haematoxylin for 6 mins and placed in running tap water for 4 mins for sample blueing. Counterstaining with Eosin Y was performed for 2 mins prior to briefly rinsing with water before dehydrating in 70%, 95%, 100%, 100% for 30 sec each and then xylene for 10 mins twice. Slides were mounted with DPX and images taken at 10X using an Olympus BX43 upright light microscope with the IC capture software (v 3.7.8.2).

Masson’s Trichrome (Abcam, AB150686) staining was carried out using the above dehydration/rehydration steps and the manufacturers protocol with several modifications: 8 mins Biebrich staining, 15 mins in phosphotungstic acid solution, 12 mins Analine blue staining and incubating for 3 mins in 1% acetic acid. Images were taken at 10X magnification.

Skeletal muscle tissue damage was assessed using a multiparametric scoring tool for assessing ischaemia severity as described previously [24]. Two blinded independent investigators scored the degree of inflammation, fibrosis, intermuscular adipocyte infiltration, necrosis, and muscle regeneration.

### Statistical Analysis

Data were tested for normality using a Shapiro-Wilk test and then assessed using either a One-way ANOVA or a Kruskal Wallis test as appropriate using Graphpad Prism (v 9.5.1). The Sidak post-hoc test was used for normally distributed data, or the Dunn’s test for non-parametric data. For all parametric two-sample analysis an unpaired Student’s T-test was used. Two-sample non-parametric ordinal data were compared using a one-way Wilcoxon test. Pearson correlation and linear regression analysis were performed between FLAIR data, and mean diffusivity or fractional anisotropy using Graphpad Prism. Statistical significance was defined at P ≤ 0.05. All data are represented as mean ± standard deviation (SD) unless stated otherwise.

All MRI data are displayed as a ratio of the ischaemic to non-ischaemic limb.

A Pearson correlation analysis was carried out between the FLAIR intensity values and the fractional anisotropy (FA) and mean diffusivity (MD) to determine the link between inflammation and the muscle fibre integrity. The Pearson correlation was carried out using Graphpad Prism (v. 9.5.1).

## Results

### 3D-GRE shows significant increases in muscle volume on Day +3 post-HLI

HLI is characterised by a loss in muscle mass due to ischaemia-induced necrosis and inflammation [10]. To determine whether a change in muscle volume in the ischaemic limb post-surgery was detectable by MRI, we used 3D-GRE scans. Results show that the ischaemic limb muscle volume was significantly increased on Day +3 compared to Day −3 (P = 0.0002), but returned to baseline levels by Day +7 (Figure 2A+B). The lack of difference in the volume at Day +7 agrees with the absence of muscle weight difference noted upon dissecting the muscles (Figure 2C+D).

**Figure 2.**
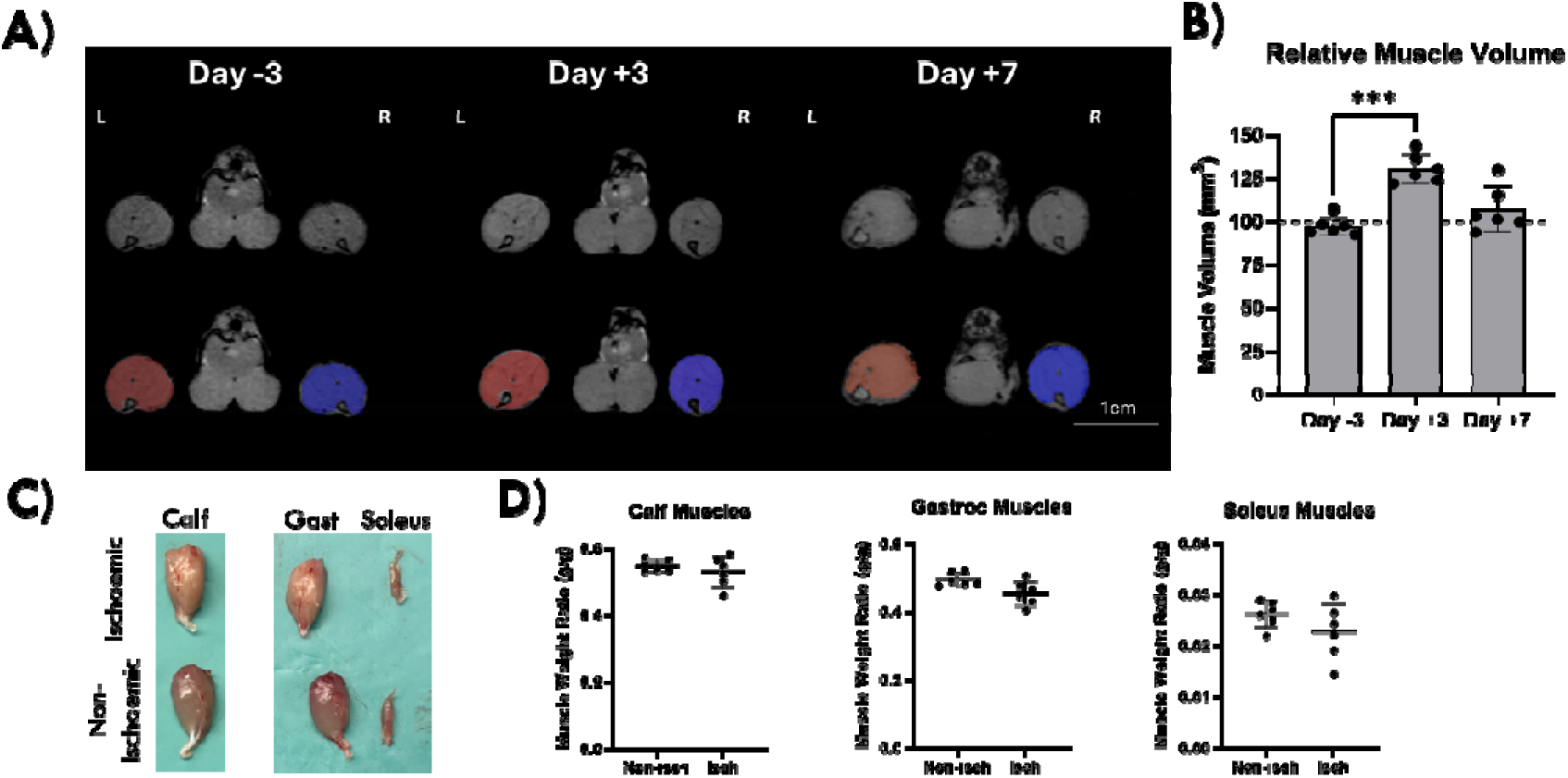
3D-GRE Muscle Volume Analysis. A) Representative anatomical 3D-GRE images of muscles. B) Volumes of the calf muscles were calculated as a ratio of the non-ischaemic limb. There was a significant increase in muscle mass on Day +3 after HLI vs Day −3 (P = 0.0002). This was restored to baseline levels by Day +7. C) Representative dissections of lower limb muscles showing significant pigmentation loss in ischaemic muscles. D) Wet weights of calf muscles of the hindlimb. No significant difference in muscle weights between ischaemic + non-ischaemic at Day +7. Gast = Gastrocnemius. N = 6. Mean±SD. Scale Bar = 1 cm. ***<0.001.

### Significant increases in FLAIR measured inflammation at Day +3 and +7

A notable hallmark of HLI is the associated inflammation that occurs, particularly at early timepoints [10,25]. FLAIR imaging was used to quantify water content providing an indirect measure of tissue inflammation [26,27]. Therefore, we generated volumes of interest around the calf and estimated the relative pixel intensity on FLAIR scans as a measure of the degree of inflammation. This was then calculated as a ratio of the ischaemic vs non-ischaemic limb. We noted a significant increase in the degree of inflammation in the ischaemic limb vs the non-ischaemic limb on Day +3 and +7 vs Day −3 (P < 0.0001 and P < 0.0001, respectively) (Figure 3A+B). Histological analysis of the ischaemic limb at Day +7 confirmed the presence of severe inflammation, characterised by an accumulation of leukocytes scattered across large areas of the muscle, accompanied by a significant loss of muscle fibre integrity (fibre degeneration) as well as large areas of muscle fibre necrosis (purple-coloured muscle fibres in Masson Trichrome staining) (Figure 3C-D). At this early timepoint, there was a mild deposition of intermuscular collagen fibrillar material (blue deposits in Masson’s Trichrome staining), but there was no presence of intermuscular adipocyte infiltration, which is more obvious in later timepoints [12]. The cumulative ischaemic score (cISS) was calculated for each animal by adding up the individual scores (Figure 3E+F), with a total cISS = 10.5 noted in the ischaemic limb, indicating severe ischaemic damage [24]. In contrast, the non-ischaemic limb possessed very mild inflammatory and fibrotic scores (cISS = 1.5), as previously observed at this timepoint [25,28].

**Figure 3.**
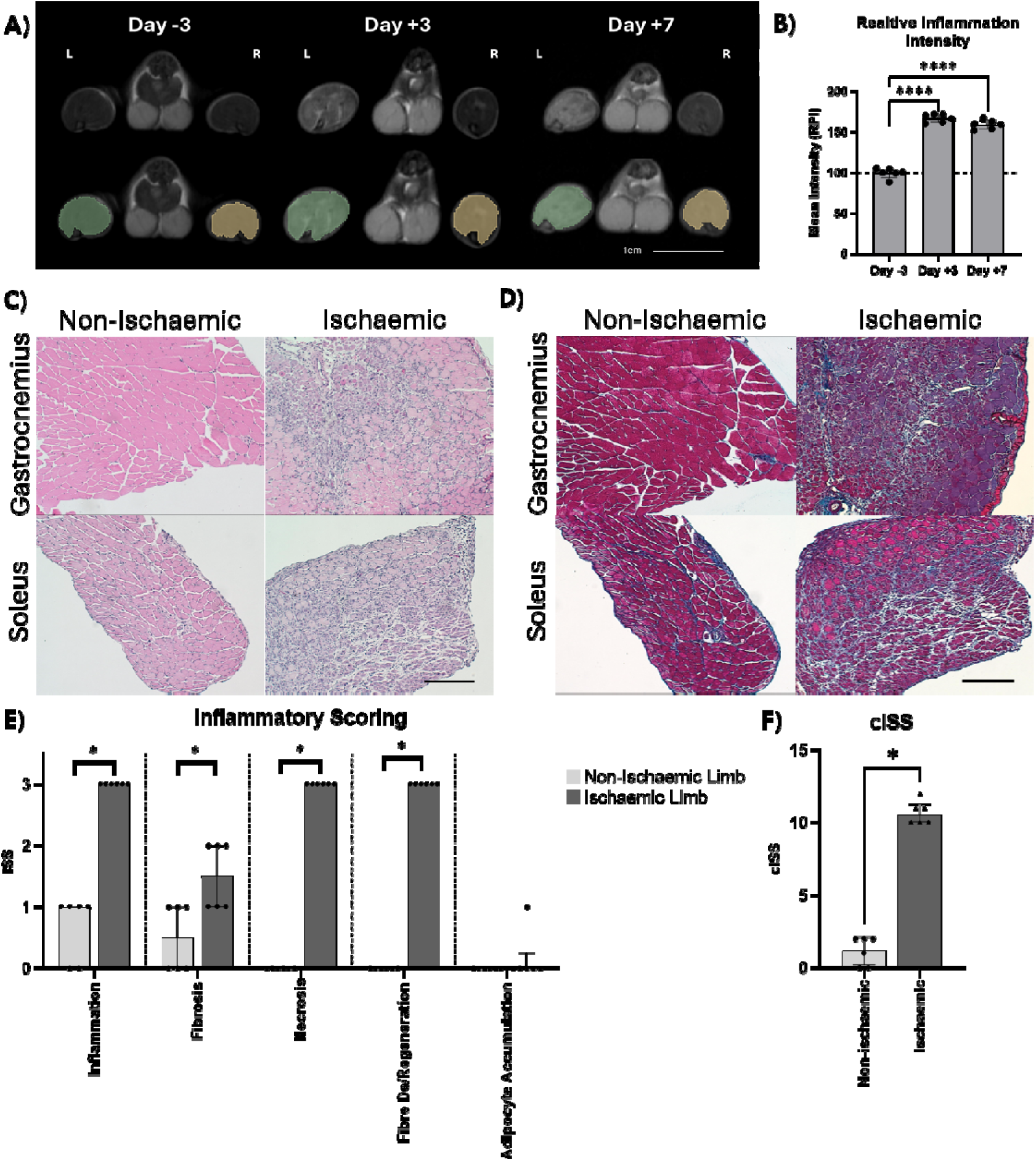
Significant inflammation observed in ischaemic muscles post HLI. A) Representative FLAIR images of inflammation in leg muscles. Scale Bar = 1 cm. B) Muscle inflammation was measured using volumes of interest of the muscle distal to the knee joint and then calculating the average pixel intensity within the volume of interest. The pixel intensity was then taken as a ratio of the ischaemic to non-ischaemic limb. There was a significant increase in muscle water content, indicative of inflammation, on Day +3 and +7 after HLI vs Day –3 (P < 0.0001 and < 0.0001, respectively). Mean ± SD. Representative H&E (C) and Masson’s Trichrome (D) histological images. Scale bar = 200 μm. E + F) Individual and cumulative ischaemia score indicates significant increases in inflammation, fibrosis, necrosis, and fibre degeneration in ischaemic muscles vs non-ischaemic control limbs. N = 6, median ± interquartile Range. * < 0.05, **** < 0.0001. RPI = Relative pixel intensity.

HLI is a complex and severe condition that results in many pathological changes including tissue necrosis. With the MRI’s ability to detect white tract destruction in the brain, we wanted to discern its ability to identify changes in muscle fibre integrity. Using DW-EPI images, we calculated the FA and MD values in three regions of the calf muscles using two consecutive slices per region, beginning just below the knee joint (superior, middle, and inferior) (Figure 4A-D). The FA and MD were then calculated as a ratio of the ischaemic vs non-ischaemic limb (Figure 4E+F). We found a significant decrease in the FA values on Day +3 after surgery, followed by partial restoration on Day +7. There was only a significant change in the MD values in the mid-calf, with an increase on Day +3 and subsequent decrease on Day +7 (P = 0.03 and 0.01, respectively).

**Figure 4.**
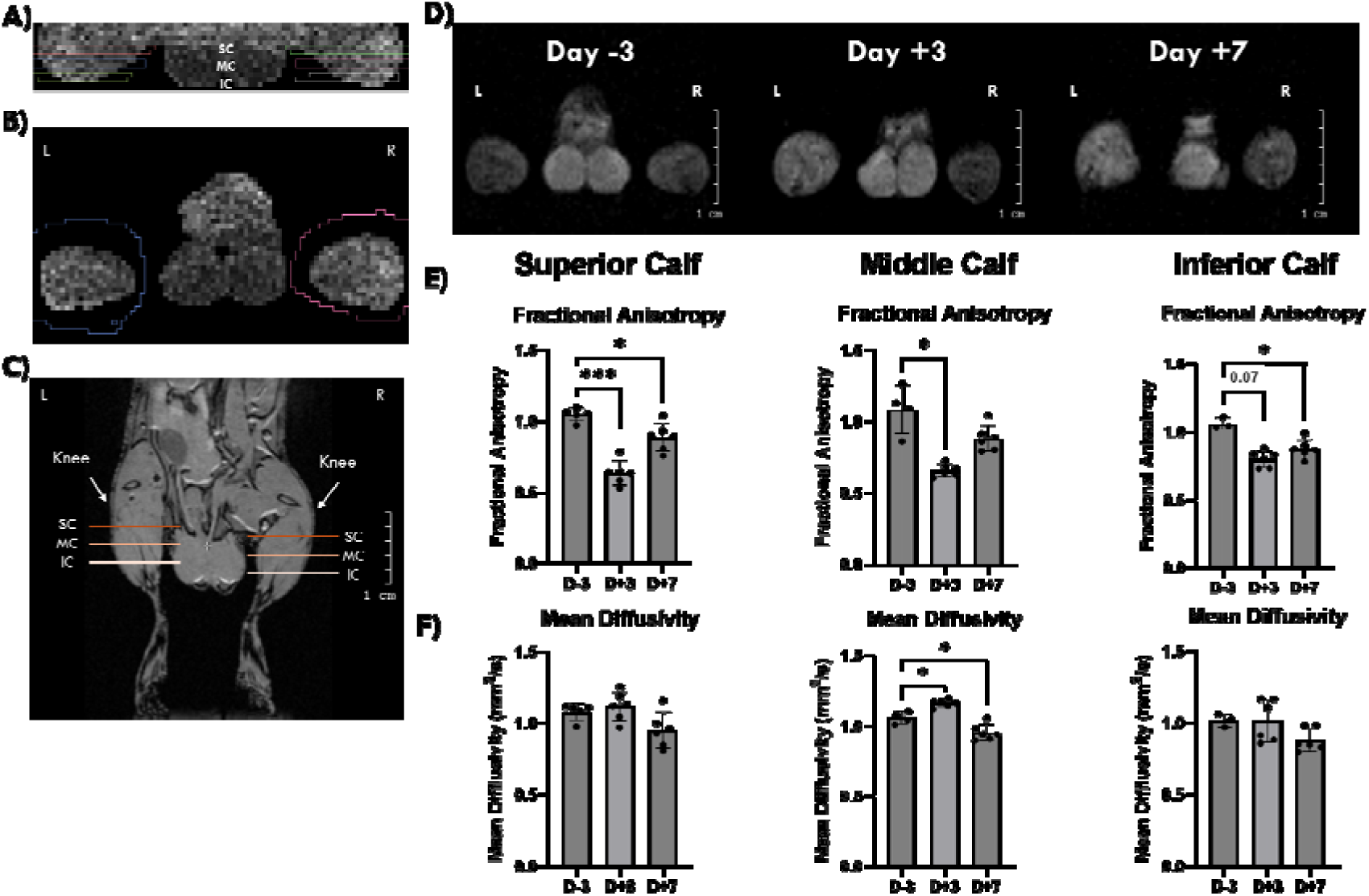
Significant decrease in muscle fibre organisation after HLI. A-C) Representative volumes of interest. D) Representative DW-EPI image of the middle calf at different timepoints. E) Fractional anisotropy of calf muscles indicating a significant decrease in muscle fibre linearity on Day +3 and +7 after surgery throughout the calf muscles. F) Significant increase in water movement noted by Day +3, likely due to oedema, and significantly restricted water movement, likely associated with immune cell infiltration, on Day +7 in the middle calf region. Both the fractional anisotropy and mean diffusivity were calculated as a ratio of the ischaemic to the non-ischaemic limb. N = 3-6. Mean±SD. * <0.05, *** <0.001. IC = Inferior Calf, MC = Middle Calf, SC = Superior Calf.

The diffusion data were then correlated with the FLAIR water content data to investigate the association between inflammation and tissue damage. There was a significant strong negative correlation between the FLAIR and FA values at each level of the calf muscles (superior (P < 0.0001, r = −0.87), middle (P < 0.0001, r = −0.87), inferior (P = 0.0017, r = −0.74)) (Figure 5A+B), and a significant moderate correlation between the MD and FLAIR values at the mid-calf region only (P = 0.03, r = 0.54).

**Figure 5.**
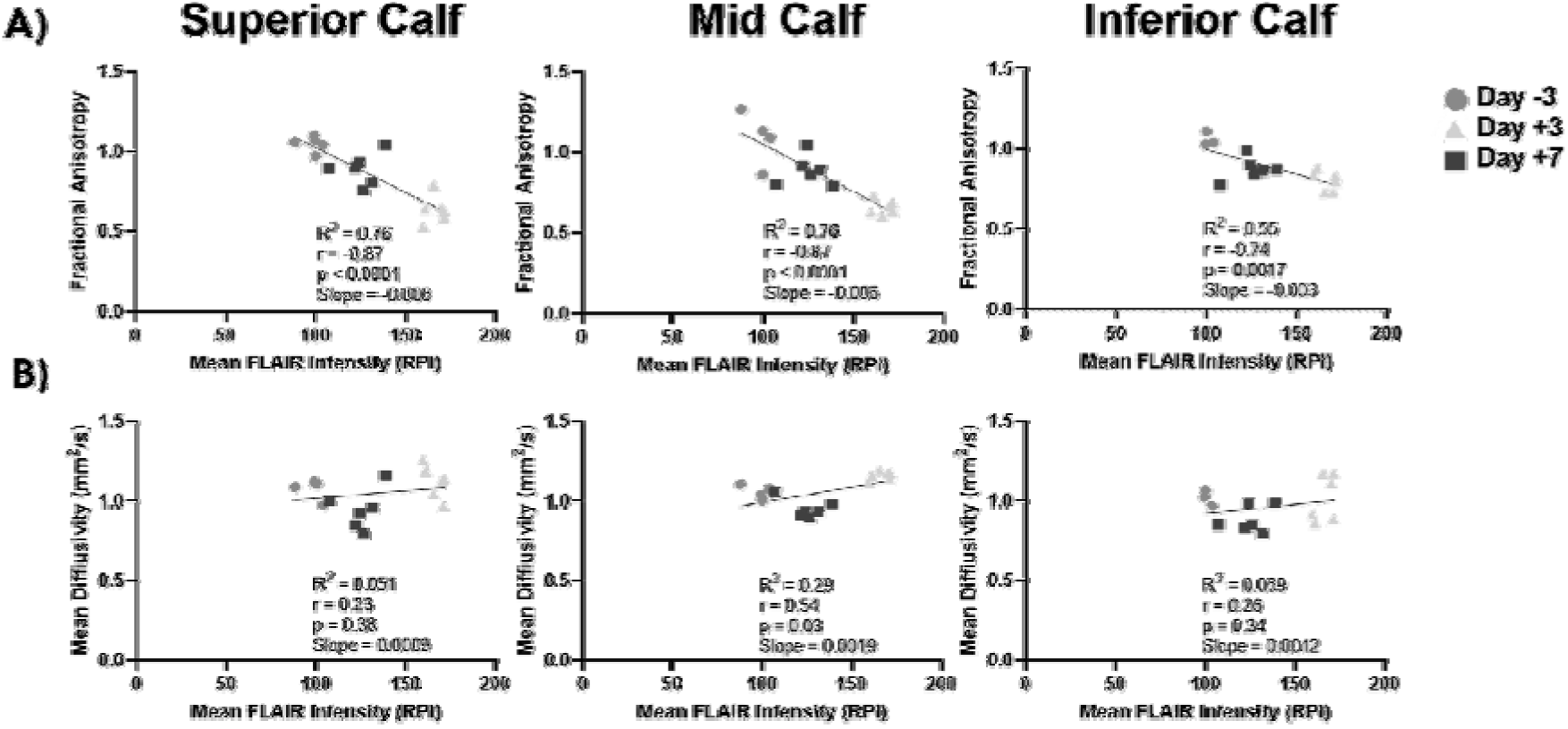
Pearson correlation of FLAIR and DW-EPI. A) The correlation between FLAIR and fractional anisotropy shows that muscle fibre integrity has a significant and strong negative correlation with water infiltration, as detected by FLAIR. This data suggests that the muscle fibre integrity decreased in tandem with inflammation. B) The water content (FLAIR) only significantly correlated with the mean diffusivity in the middle calf region, suggesting that the ratio of the cell and water content was mostly unchanged in the calf. RPI = Relative pixel intensity.

### Significant decrease in arterial tree length and perfusion after HLI

Restoring blood flow post-ischaemia is a critical step towards functional limb recovery. Thus, identifying vascular remodelling via MRI along with restored blood perfusion are crucial metrics to determine the ability of the body to restore nutrients to the nearby ischaemic tissue. Using 3D-TOF imaging, we validated successful surgical ischaemia with a corresponding significant decrease in the max length of vessels in the ischaemic limb vs the non-ischaemic limb at Day +3 and +7 vs −3 (P < 0.0001 and 0.0001, respectively) (Figure 6A-C). By Day +7, there was no substantial increase in max arterial length identified. Using ASL imaging, we noted a corresponding significant decrease in the perfusion in the ischaemic limb by Day +3 and Day +7 compared to pre-surgery values (P = 0.03 and 0.04, respectively) (Figure 6D-F).

**Figure 6.**
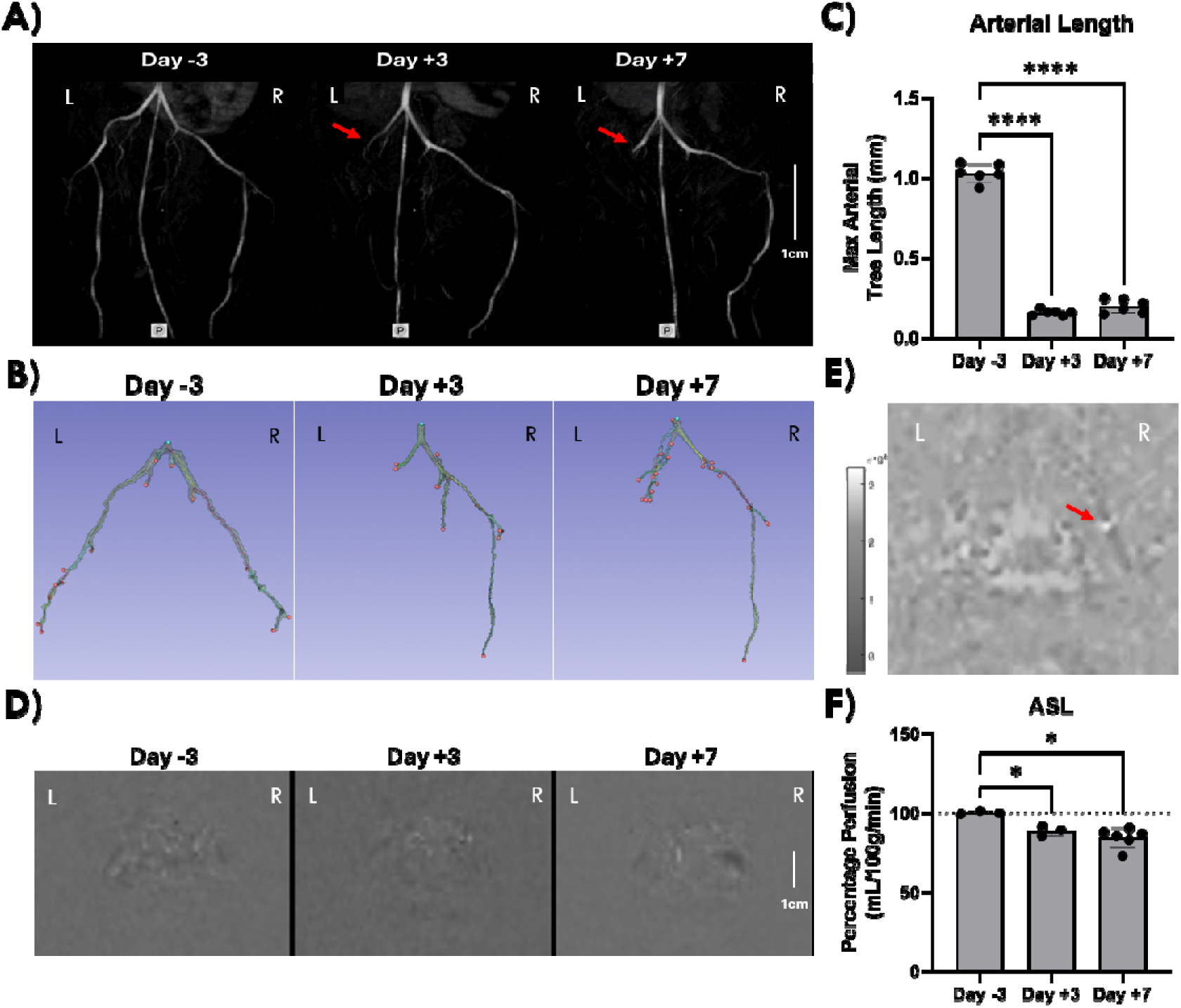
Limb Vascular Deterioration. A) Representative TOF images of vessels in the hindlimb. MRI can identify the vasculature of the lower limb and the site of surgical arterial transection (red arrow). Scale bar = 1cm. B) Model of the surface profiling at each timepoint. C) Graph of the longest combined arterial length beginning from the descending aorta-common femoral branch junction to the furthest arterial branch in each limb. The arterial length was then calculated as a ratio of the ischaemic to non-ischaemic limb. Our findings show a significant decrease in the longest combined length of artery in the ischaemic limb after HLI surgery compared to Day-3 (P < 0.0001). N = 6. D) Representative perfusion images at Day −3, +3, and +7. E) Processed image of hindlimb for perfusion modelling. The red arrow indicates a region of high perfusion in the non-ischaemic limb, with no corresponding high perfusion in the ischaemic limb. F) Quantification of the relative perfusion in ischaemic vs non-ischaemic limbs showing a significant decrease in perfusion after HLI surgery. N = 3-6. * = 0.05. **** <0.0001.

## Discussion

Here, we developed an MRI multiparametric approach to characterise the pathological cascade following HLI, encompassing tissue structure, vascular responses, and mechanisms underlying recovery. Overall, this study demonstrates the potential of MRI to longitudinally monitor recovery in the HLI mouse model, capturing the muscular, inflammatory, and vascular responses through the combined use of 3D-GRE, ASL, 3D-TOF, DW-EPI and FLAIR imaging.

Patients with CLTI exhibit substantial loss of muscle mass and function [29], a phenotype that our lab and others have also observed in the HLI mouse model [12,25]. This decline is likely attributable to impaired nutrient delivery, ischaemia-induced tissue necrosis, and reduced mechanical loading, which collectively promote muscle atrophy and structural deterioration. Here, we have shown that MRI can be used to track changes in muscle volume across time. At an early timepoint of Day +3 post-HLI we identified a significant increase in muscle mass using volumetric analysis of MRI images (Figure 2). This increase in volume is likely due to the oedema associated with the ischaemia induced inflammation, which was corroborated in the FLAIR imaging [30]. We predict this muscle mass will decrease at later timepoints as shown by Evans et al. [25], as suggested by the drop in muscle volume on Day +7. While data by Scalabrin et al [31] noted a significant decrease in gastrocnemius and soleus weights at Day 7 post-HLI, our data did not find significantly differences. The difference is likely attributed to Scalabrin et al transecting both the femoral artery and vein, causing a more severe phenotype. Sustained inflammation is a key hallmark of limb ischaemia and is known to contribute negatively to tissue remodelling and functional recovery. The hypoxic and inflammatory environment act as potent chemoattractant stimuli, promoting rapid immune cell infiltration while contributing to extensive tissue necrosis and oedema. Our histological data similarly shows significant immune cell infiltration and tissue necrosis at Day +7 (Figure 3). MRI analysis demonstrated water content changes, a surrogate marker of inflammation, and demonstrated significant oedema on both Day +3 and +7. In addition, at Day +7 histological assessment revealed early muscle remodelling, characterised by mild collagen deposition and large areas of muscle fibre degeneration, although extensive collagen accumulation and adipocyte infiltration were not yet evident. The absence of adipocyte accumulation suggests that this represents an early remodelling stage prior to the development of the more advanced fibrotic and adipogenic phenotype observed at later timepoints [12,13,24,32]. Our DW-EPI data shows that there was significant reduction in muscle fibre integrity post-HLI surgery at Day +3, which begins to recover by Day +7 (Figure 4), which significantly correlated with the extent of oedema as evaluated by the FLAIR imaging (Figure 5). Similar correlations have been observed between T2 relaxation times and % of muscle damage determined using histology [13]. Overall, MRI can reliably detect early muscle remodelling changes longitudinally including inflammation and muscle integrity, using changes to water influx and muscle fibre integrity.

While LDI is the gold standard technique to measure blood flow recovery post-HLI, MRI can visualise and functionally validate the surgical ligation. This can be done by directly imaging the vessels while also comparing perfusion between the ischaemic and non-ischaemic limbs. MRI imaging confirmed successful induction of the HLI model, demonstrated by a significant reduction in perfusion in the ischaemic limb with ASL imaging at Day +3 and +7. This finding was further supported by 3D-TOF imaging, which revealed a significant reduction in arterial tree length (Figure 6). The decrease in perfusion is similar to that observed in HLI studies that use LDI [33–35]. However, ASL demonstrated a decrease in perfusion of only ∼15% of the ischaemic vs non-ischaemic control after HLI surgery. In contrast, LDI typically demonstrates a drop in perfusion of ∼80% perfusion after HLI, suggesting a higher sensitivity to detecting changes in perfusion. A study by Bryant et al. [18] showed potential for dynamic contrast imaging for detecting perfusion changes in inflammation; therefore, future work can assess whether dynamic contrast-enhanced MRI provides superior sensitivity compared to ASL in HLI.

The main limitations of this work are the long imaging times required (e.g. 2 hrs per animal for all the scan modalities), complexity of the analysis, expertise, and cost of an MRI machine. Despite this, multimodal MRI has the advantage of providing comprehensive information on muscle structure, inflammation, and vascular remodelling. It facilitates longitudinal analysis of multiple parameters in vivo without the need for animal sacrifice. Nevertheless, LDI may remain a more practical approach for longitudinal monitoring of perfusion recovery due to its simplicity and resolution, and perhaps can be applied in parallel. Furthermore, our data demonstrates low variability in muscle volume, inflammation, muscle fibre integrity, and vascular monitoring data. Therefore, even with low animal numbers, significant changes were identified between the ischaemic and non-ischaemic limbs. Finally, we observed no significant functional and anatomical recovery by Day +3 and +7, which is likely due to endogenous vascular recovery beginning at later timepoints following HLI induction [13,36]. Therefore, further studies incorporating later timepoints will be required to fully characterise the temporal progression of the endogenous HLI recovery process. Finally, this work did not investigate a potential therapeutic to discern its therapeutic mechanism of recovery. To demonstrate the maximum potential of longitudinal MRI, future work should investigate novel therapeutics in the HLI model.

In the current study we demonstrate the feasibility of using MRI for multimodal in vivo sequential analysis of the response to limb ischemia. This report also lays the foundation to apply mutli-modal MRI imaging to examine the recovery of HLI mouse models in response to novel therapeutics. MRI can provide invaluable information on the mechanisms of action of new therapies, thereby accelerating their translation to the clinic. Subsequent investigations should examine recovery at later timepoints, as many HLI studies extend follow-up to 21 or 28 days post-ischaemia. Incorporating later imaging timepoints will provide invaluable information into the endogenous recovery process following HLI induction, including the resolution of inflammation, vascular remodelling, and restoration of muscle fibre integrity to determine whether these processes are accelerated by therapeutic interventions.

### Conclusion

This work demonstrates the potential of multimodal MRI as a longitudinal tool to monitor recovery following HLI induction in vivo, enabling detection of changes in muscle fibre integrity, vascular remodelling, and inflammation over time. Future work can apply MRI technology to help elucidate the mechanism of action of therapeutics to better facilitate their clinical translation.

## Acknowledgements

We would like to thank Gabor Mosaic from Mediso Ltd (Hungary) for his technical input on the MRI and to CÚRAM, Research Ireland Centre for Medical Devices, University of Galway, Galway, Ireland for the use of the MRI machine. Figures were made with the help of Biorender.

## Author Contribution

C.J.L. = Conceptualisation, data acquisition, data analysis, manuscript writing.

M.N.D. = Data acquisition.

X.Z.C. = Data acquisition.

C.A.L. = Data acquisition and manuscript review.

C.S.N. = Data acquisition and manuscript review.

T.O.B. = Conceptualisation, funding acquisition, manuscript review.

N.C. = Conceptualisation, data analysis, manuscript writing, and review.

## Abbreviations

cISS: Cumulative Ischaemia Severity Score
CLTI: Chronic Limb Threatening Ischaemia
DW-EPI: Diffusion Weighted Echo Planar Imaging
FA: Fractional Anisotropy
FLAIR: Fluid-Attenuated Inversion Recovery
Gast: Gastrocnemius
GRE: Gradient Echo
HLI: Hindlimb Ischaemia
IC: Inferior Calf
MC: Middle Calf
MD: Mean diffusivity
MRI: Magnetic Resonance Imaging
SC: Superior Calf
TOF: Time of Flight

